# Developing Future Biologists: creating and assessing a portable short course to engage underrepresented undergraduate students in developmental biology

**DOI:** 10.1101/467092

**Authors:** Justine M. Pinskey, Eden A. Dulka, Andrea I. Ramos, Martha L. Echevarría-Andino, David S. Lorberbaum, Brandon S. Carpenter, Jorge Y. Martinez-Marquez, Breane G. Budaitis, Emily M. Holloway, Samhitha Raj, Alana M. Chin, Edu Suarez, Laura A. Buttitta, Deb L. Gumucio, Deneen M. Wellik, Ryan Insolera, Leilani Marty-Santos, Benjamin L. Allen, Scott Barolo

## Abstract

Many barriers discourage underrepresented students from pursuing science careers. To access graduate education, undergraduate students must first gain exposure to a particular subject and subsequently accumulate related coursework and research experience. Many underrepresented students lack exposure to developmental biology due to limited undergraduate course offerings and finite resources at smaller institutions. To address this disparity, a group of University of Michigan graduate students and postdoctoral fellows created a portable short course focusing on developmental biology, titled “Developing Future Biologists” (DFB). This weeklong educational initiative provides hands-on laboratory sessions, interactive lectures, and professional development workshops to teach students about developmental biology and increase awareness of scientific career options. To evaluate course effectiveness, we developed a pre-post assessment, incorporating main ideas from the BioCore Guide. Student understanding of basic concepts and perceived experience in developmental biology increased in DFB participants, despite the abbreviated nature of the course. Here, we provide all course materials and an in-depth analysis of the assessment we created. The DFB portable short course model is an easily adaptable tool that connects undergraduate students with opportunities for advanced study and lowers barriers for underrepresented students in science, technology, engineering, and mathematics.

## Introduction

Similar to other science, technology, engineering, and mathematics (STEM) fields, developmental biology trainees do not proportionally represent the diversity of our nation (NSF (National Science Foundation), 2015). Many barriers contribute to this lack of diversity, including limited opportunities to partake in relevant science coursework and gain research experience (Hurtado, Newman, Tran, & Chang, 2010). It is widely accepted that increased diversity enhances graduate student training and development through the integration of a variety of cultural perspectives (Aguilera, 2012). For outstanding students from all backgrounds to join the developmental biology community, however, they need to be made aware of opportunities in science and develop a passion for this exciting field.

To address this issue, a team of University of Michigan graduate students and postdoctoral fellows created Developing Future Biologists (DFB), an educational initiative designed to lower the cultural barriers to graduate education, increase awareness of science careers, and teach students core concepts of developmental biology. The program centers on a weeklong short course that includes developmental biology instruction, hands-on laboratory exercises, professional development activities, and networking sessions. A main focus of the course is to incorporate active learning strategies, which enhance student learning in STEM fields (Freeman et al., 2014; Haak, HilleRisLambers, Pitre, & Freeman, 2011). Additionally, the course aimed to assist students from a variety of backgrounds build long-term mentoring relationships with University of Michigan graduate students, postdoctoral fellows, and faculty members. Previous research suggests that similar mentoring efforts have helped students succeed in scientific endeavors (Tsui, 2007). Importantly, DFB was designed to be portable and scalable so that future iterations could be adaptable to a wide variety of subjects and locations.

In May of 2015, our team implemented the first DFB course in Ponce, Puerto Rico, where several of our instructors had completed undergraduate studies in biology and noticed a need for developmental biology instruction. Only three of the ten University of Puerto Rico (UPR) undergraduate campuses offer a developmental biology course on a regular basis, with only two of the three offering laboratory-based instruction. Therefore, after a successful pilot program focused on UPR Ponce students, we returned to Puerto Rico in 2016, opening the application to undergraduate students from all UPR campuses and providing room and board for students from other cities. To accomplish this, we compiled a variety of external and internal funding from sources such as the Society for Developmental Biology Non-SDB Educational Activities Grant, American Society for Cell Biology Committee for Postdocs and Students Outreach Grant, the Department of Cell and Developmental Biology at the University of Michigan, and the Rackham Graduate School Dean’s Strategic Initiative at the University of Michigan, among others (see Acknowledgements). In addition, we offered a local iteration of the course to underrepresented undergraduate students from the state of Michigan in 2017, allowing us to validate our assessment on a second student demographic.

Since 2010, the Vision and Change Call to Action has been instrumental in guiding innovations in biology education (AAAS, 2010). The DFB course model incorporates many aspects of Vision and Change, including integrating core concepts into the curriculum, focusing on student-centered learning through active participation, and encouraging students and professors from the University of Michigan to embrace high quality, innovative teaching methods. To gauge the effectiveness of the course and measure students’ grasp of core concepts in developmental biology, we developed and incorporated formal pre-post assessments using the BioCore guide’s interpretation of the Vision and Change core concepts in Biology (Brownell, Freeman, Wenderoth, & Crowe, 2014). To assess student attitudes, we also included a pre-post survey asking students to self-rate their experience in developmental biology, their interest in graduate school, and their awareness of science career options.

The purpose of this report is two-fold: 1) To present data addressing the effectiveness of the 2016 and 2017 DFB course iterations, and 2) To provide resources for the development of similar initiatives. Throughout the course, our goals were to build relationships with students, improve attitudes about graduate education and scientific careers, and effectively teach students the core concepts of developmental biology. Here, we present our course design, teaching materials and assessment data, demonstrating that this short course model can enhance student understanding and perceived experience in developmental biology.

## Course Development and Methods

### Instructor Selection and Preparation

Graduate student, postdoctoral fellow, and faculty instructors for the DFB course were selected eight months before the course start dates. Faculty instructors were selected based on involvement with undergraduate and graduate education and were invited to participate via email. All other instructors submitted a cover letter and curriculum vitae, and selected applicants were interviewed. Instructors were selected based on teaching experience, interest in social justice/inclusion, and ability to commit two years to the program. A two-year commitment was required to help with turnover and ensure the continuation of the initiative. Over the course of the academic year, instructors met weekly to develop the course curriculum, design lab activities, create the course applications and advertisements, design assessments, review applications, practice lab instruction, and perform other tasks related to creating this course. Meeting notes, as well as all other materials created for the course, were organized and stored on a shared drive.

### Student Applications & Selection

Approximately three months before the course start date, advertisements were sent out via email, flyers (Figure S1), the course Facebook page (https://www.facebook.com/developingfuturebiologists), and the course website (http://developingfuturebiologists.com). Applications, which were designed using Google Forms and linked through the course website, were open for one month. The course was restricted to 24 students due to equipment limitations and to maximize the personal interactions among enrolled students and DFB instructors. Students were selected for participation based on grade point average (minimum of 2.8 on a 4.0 scale) year in college, major, career goals, and reason for course interest. While senior students participated in the course, we sought to accept a larger portion of first and second year students based on previous studies demonstrating that early research experiences increase continued participation in the sciences, particularly for students from underrepresented minority groups (Nagda B.A., 1998; Rodenbusch, Hernandez, Simmons, & Dolan, 2016). Preference was given to students who communicated enthusiasm and a clear personal benefit from participation in DFB.

### Lectures and Labs

One of our main goals in developing this course was to teach students the core concepts of developmental biology. To achieve this objective, we developed interactive, discussion-based lectures and hands-on laboratories surrounding main ideas and experimental techniques in developmental biology. Each day focused on a single theme, including early embryonic development, cell signaling, gene expression, organogenesis, and development and disease. Instructional sessions were held daily from 9:00 AM to 5:00 PM, with discussion-based lectures in the morning, interactive labs in the afternoon, and occasional evening networking activities (Figure S2).

Prior research indicates that courses with hands-on activities, such as labs, increase enthusiasm and learning in students (Basey et al., 2014). Therefore, our course was designed to focus on lab activities, with discussion-based lectures serving to introduce core content that was later incorporated into laboratory material. Experienced faculty, postdocs, and graduate students led discussions, and slide presentations were combined with active-based learning to encourage student involvement. For example, one discussion session included reenactment of the Wnt cellular signaling pathway, where students acted out the functions of specific pathway components. In addition, the use of iClicker remotes allowed instructors to pose questions throughout the lecture to further engage students in the material (Caldwell, 2007; Crossgrove & Curran, 2008).

Afternoon labs served as the main hands-on component of the course, with the objective of exposing students to basic research methods and tools used in the field of developmental biology. Instruction in basic laboratory safety and record keeping was provided prior to lab participation. Due to space limitations in 2016, the course was designed to have two lab sessions covering the same material each afternoon; half of the students attended lab, the other students attended professional development sessions, and then the groups switched sessions after 90 minutes. This format was kept in 2017, as positive feedback from 2016 professional development sessions demonstrated a strong need for such instruction. Each lab session utilized materials from common model organisms, including worms, frogs, flies, chickens, and mice. To guide participation, students were provided a lab workbook containing background information, experimental protocols, questions about each specific exercise, and space for students to record their observations (Figure S3). Workbooks were not graded, and students were allowed to keep their workbooks following the conclusion of the course.

### Professional Development for Participants

The importance of mentoring in underrepresented student success is well established (Nagda B.A., 1998; Villarejo, Barlow, Kogan, Veazey, & Sweeney, 2008; Whittaker & Montgomery, 2012). To facilitate mentoring relationships between DFB instructors and participants, students were split into groups and assigned two team leaders from the University of Michigan. The team leaders mentored their assigned students both during the course and after its conclusion. To encourage bonding within these teams, friendly competitions including questions about course content and lab-based challenges were held throughout the week for small prizes. In addition to the team assignments, networking activities at which students could informally interact with their assigned mentors as well as other instructors were held. These included an ice cream social, a bowling night, dinner with additional faculty from outside of the course, and an end-of-course networking dinner. These events created a welcoming environment for undergraduates to speak with graduate students, postdocs, and faculty, encouraging the formation of meaningful mentoring relationships.

We further aimed to help students envision themselves as researchers and learn about career opportunities by incorporating a series of career development and informational sessions. Topics included *curriculum vitae* review, effective networking skills, interview skills, presentation skills, and program-based opportunities offered at institutions like the University of Michigan (Figure S2). Additionally, career panels were incorporated into the course: one with current graduate students, and one with and faculty/postdoctoral instructors. The panels began with introductory statements from each member describing their personal scientific career paths, and proceeded with questions focused on careers in research. To allow students to gain further perspective on the types of projects and scope of work done by graduate students, the graduate student instructors gave short research talks about their specific projects daily during lunch.

### Assessment tools and statistical analyses

Previous reports have outlined the major concepts in biology, including an adaptation for developmental biology in particular (Brownell et al., 2014; Rossi et al., 2013). Using these as a guide, we created pre-post assessments to evaluate student understanding of core concepts in developmental biology as well as lab techniques used throughout the course (Figure 1). Questions corresponded to each of the major topics of the course, each with five possible answers (Figures S4a: questions 6-29, S4b: questions 6-35). Question order was randomized from pre- to post-test to avoid memorization of the questions. A score of 0 and 1 was assigned for each incorrect and correct answer, respectively. Additionally, five items on the pre-post examination were included to ascertain students’ general perspectives about scientific careers and developmental biology (Figures S4a,b, questions 1-5). Each answer was given a value of 0-4 with 0 corresponding to least familiar/positive about the question topic and 4 being most familiar/positive about the question topic. All values are reported as mean ± standard deviation. Normality of the data were tested with D’Agostino & Pearson normality tests and statistical comparisons between pre-test and post-test results were made using paired two-tailed *t*-tests, Wilcoxon matched-pairs tests, or one-way ANOVA with Sidak’s *post hoc* test as dictated by design and data distribution. Importantly, all DFB instructors completed a training certification for research in human subjects through the Program for Education and Evaluation in Responsible Research and Scholarship (PEERRS) at the University of Michigan, and this study was formally exempt from ongoing Institutional Review Board (IRB) review. Furthermore, participants were provided the option of signing a release form for photography and videography taken during the course.

**Figure 1.**
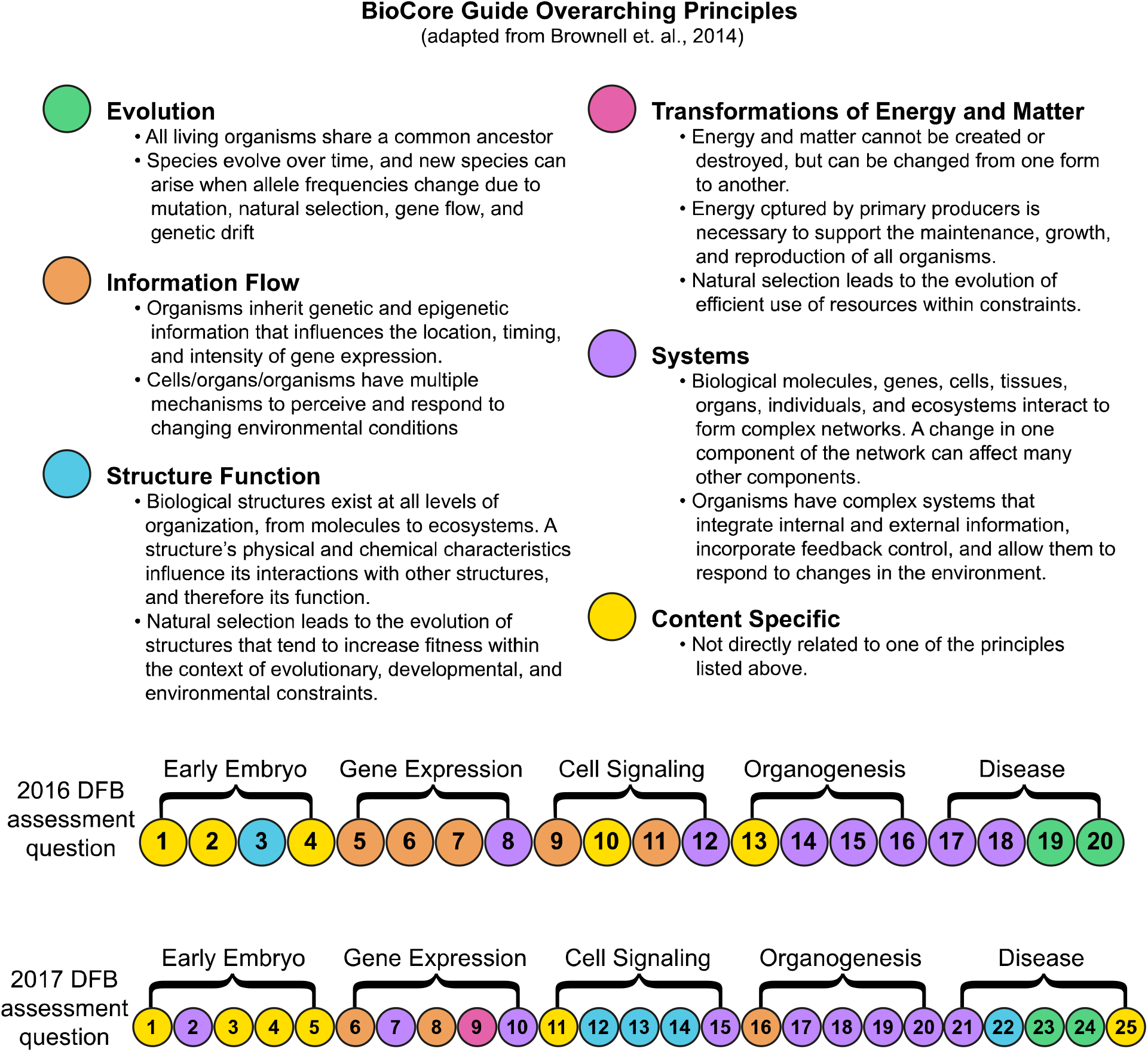
The DFB pre-post assessment covers the core concepts of developmental biology. The core concepts in Biology were defined by Vision and Change (AAAS, 2010) and further adapted into overarching principles by the BioCore Guide (Brownell et. al. 2014). Each question within the DFB pre-post assessment from 2016 (top row) or 2017 (bottom row) either relates directly to an overarching principle or relates to specific content covered in lecture or lab, as indicated by color coordination. See Supplemental Figures 6 and 8 for full assessments.

Feedback for the course was collected on a combination of iClicker data, pre-post tests, and written feedback surveys. Student feedback was collected for each lecture and lab session at its conclusion, via a standard series of iClicker questions and a written feedback survey (Figure S5, data not shown). To gain qualitative insight into the students’ experiences, we also encouraged open-ended reflections on the back of the course evaluations.

## Results

### Student understanding of core concepts improved over the course of the week

To measure understanding of core concepts in the field of developmental biology, we analyzed scores from pre- and post-test questions addressing our five content areas (Figure 2). In 2016, students’ background knowledge varied widely, with pre-assessment scores ranging from 17-50% (4 to 12 correct out of 24) and average score of 33% (8 correct out of 24). Student background knowledge prior to the course in 2017 was similar, with pre-assessment scores ranging from 13-60% (4 to 18 correct out of 30) and average score of 36% (11 correct out of 30). Excitingly, for both course iterations, cumulative student understanding of the core concepts in developmental biology improved (Figure 2A). Post-test scores rose from an average of 33% to 57% (raw score of 7.9±2.7 to 13.8±3.7, p<0.0001) in 2016 and from an average of 36% to 66% (raw score of 10.9±3.7 to 19.9±4.6, p<0.0001) in 2017. Overall, 22 out of 24 students in 2016 and all 15 students in 2017 improved their scores on the post-assessment, with increases ranging from an additional 1 to 10 points out of 24 in 2016, and 3 to 22 out of 30 2017 (Figure 2B). In 2016, a single student’s test score decreased (from 11 to 8 of 24 points), and one did not change (7 of 24 points on both assessments). The average percent improvement per individual was 24% in 2016 and 30% in 2017. Accordingly, the distribution of test scores shifted to more positive values on the post-test (Figure 2C-D). Together, these results suggest an improved overall understanding of core developmental biology concepts.

**Figure 2.**
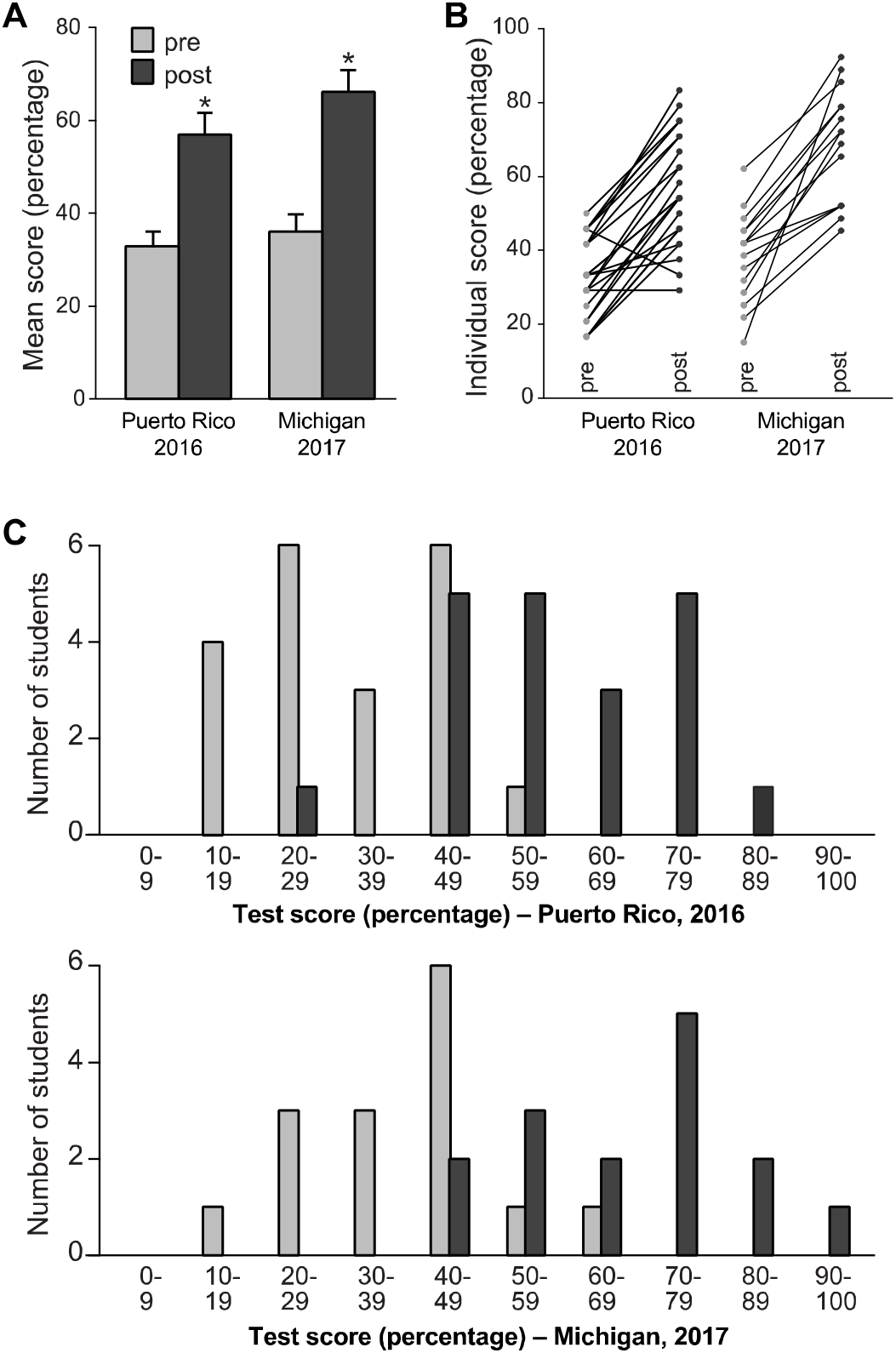
Student understanding of developmental biology core concepts improved as a result of DFB instruction. (A) Aggregate test scores on pre- and post-assessment. Data reported as mean ± SD. *p<0.0001 for both years, using a paired two-tailed Student’s t test. Score ranges: 2016, pre-assessment: 4 to 12 points out of 24 (17% to 50%; median 33%), post assessment: 7 to 20 points (29 to 83%; median 54%). 2017, pre-assessment: 4 to 18 points out of 30 (13% to 60%; median 40%), post assessment: 13 to 27 points (43% to 90%; median 70%).(B) Individual student pre- and post-assessment scores. Black lines connect each individual’s pre and post score. (C-D) Range of scores on the pre (light gray) and post (dark gray) assessments for each year.

In addition to the overall increase in test score, it was important to investigate whether a net improvement existed for each of the individual concept areas. Each test was divided into an equal number of questions assigned to each of the five concepts and a section on lab based techniques, with question order randomized. To analyze content-specific changes, questions were sorted and changes in pre-post scores were examined for each concept section (Figure 3). Understanding of all concepts improved during both iterations of the course (two-tailed t-test, p<0.05), with some concepts improving more than others. In 2016, organogenesis improved the most (40% average increase in score for that concept), followed by early embryo (32%), techniques (27%), development & disease and cell signaling (18% each), and finally gene expression (10%). In 2017, questions covering techniques improved the most (41%), followed by early embryo (40%), organogenesis (31%), development & disease (25%), cell signaling (24%), and finally gene expression (19%). These results suggest that intervention was successful for student learning each day of the course.

**Figure 3.**
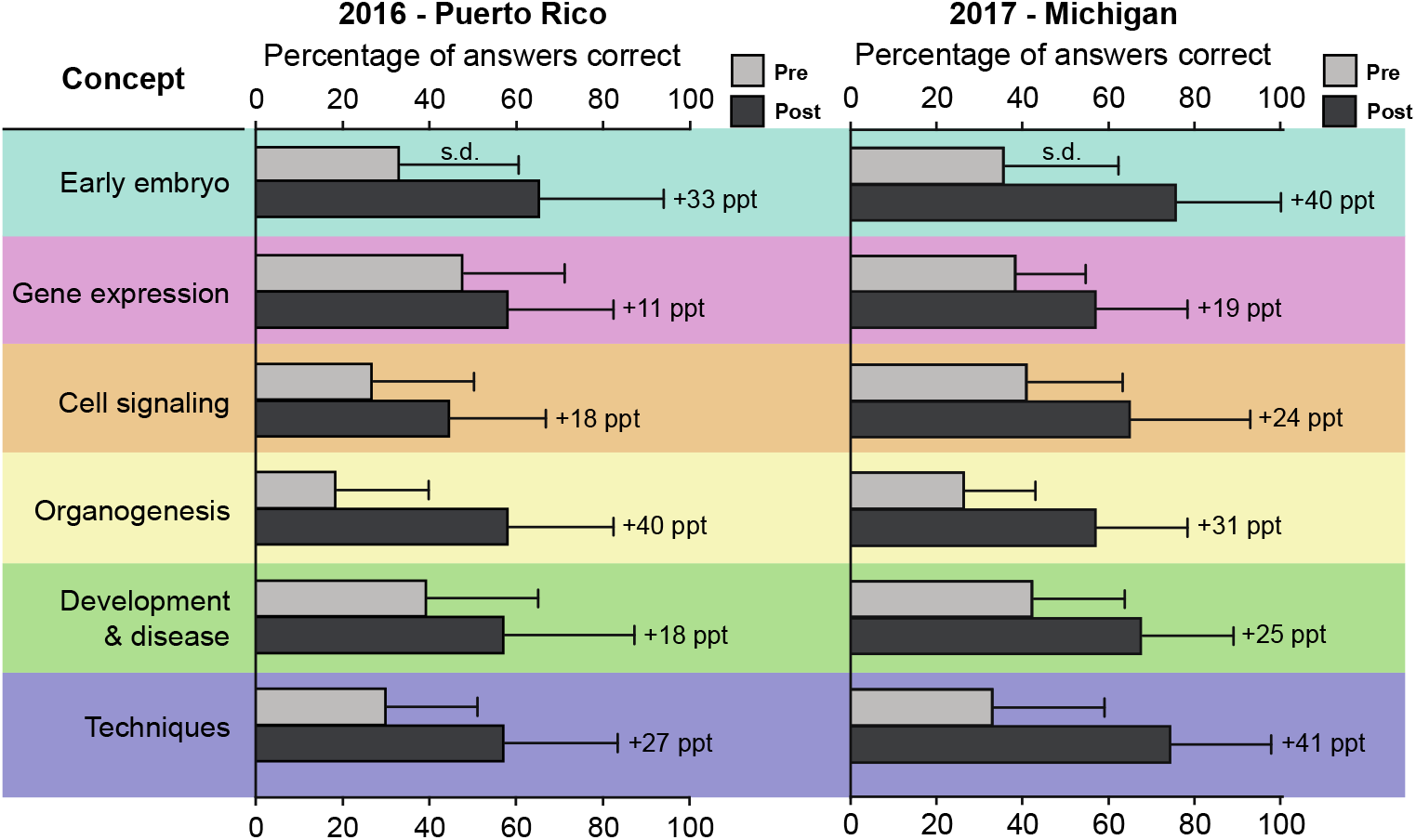
Student understanding of developmental biology improved within each conceptual area of instruction. The percentage of correct answers on the pre- and post-tests increased for all core concepts in both 2016 (left) and 2017 (right). The percentage of students who answered correctly on the pre (light grey) and post (dark grey) tests are displayed for each concept (4 questions per concept). Each concept showed significant improvement (two-tailed t-test, p<0.05). Average percent increase for each concept listed to the right of post-test bars (ppt = percentage point).

Although scores improved for the test as a whole as well as for each individual concept area, post-assessment score in 2016 was still only 57%. While conducting our initial analysis of the 2016 data, we observed that some items on the 2016 assessment were simply not covered during the course. In some cases, instructors changed their lecture content after the assessments had been designed, whereas in other cases, instructors simply ran out of time to discuss all of their material. To address this consideration, we asked instructors to self-rate coverage of each assessment item relevant to their topic on a numerical scale. After receiving all instructor analyses, questions were divided into two equally-sized groups, labeled “less instructor coverage” and “more instructor coverage”. As predicted, the topics that were covered more in depth or by multiple instructors had better learning outcomes than those topics that were not as well-discussed (Figure 4). We found that, while the “more covered” portion and the “whole” test improved from pre to post, there was no improvement in the “less covered” group of questions (percent correct mean±SD, less covered pre 37.9±14.7, post 46.5±15.5, more covered pre 27.8±16.2, post 68.4±20.0, whole test pre 32.8±11.1, post 56.9±15.2, p=0.162 less covered, p<0.0001 more covered and whole test, one-way repeated measures ANOVA/Sidak). These results demonstrate that in 2016, students improved to a greater extent on the material that the course rigorously covered than on material that was less-discussed. These findings were taken into consideration during planning of the 2017 course, leading to improvement of post-assessment scores from an average of 57% in 2016 to an average of 66% in 2017.

**Figure 4.**
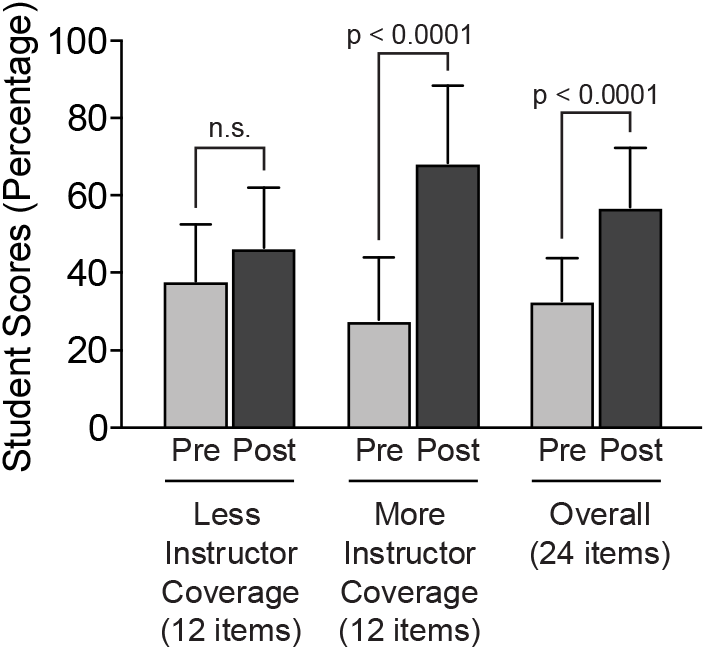
2016 Student performance correlates with increased instructor coverage. Item coverage scores were collected from instructors, and test items were divided into two equal groups reflecting less instructor coverage (less than 60% instructor coverage score), and more instructor coverage (greater than 70% instructor coverage score). Student scores by item coverage changed based on instructor coverage scores (one-way repeated-measures ANOVA, p<0.0001). Post hoc analysis results suggest students scored similarly on pre and post items for which less instruction was given (Sidak p=0.1620), while scores improved on items for which more instruction was given (Sidak p<0.0001) and for the test overall (Sidak p<0.0001).

### Item analysis provided recommendations for assessment improvement

To measure the validity and effectiveness of our assessments, we performed item analyses for difficulty for both the 2016 and 2017 pre-post exams (Tables 1 and 2). Item difficulty measures the proportion of students who answer an item correctly (Allen & Yen, 2002). Therefore, a higher difficulty score signifies that an item was “easier”, because a higher percentage of students answered that item correctly. Importantly, overall difficulty scores increased from pre to post for both years, indicating that more students answered post-test items correctly following instruction (Tables 1 and 2). In 2016, 5 out of 24 items on the pre-assessment (21%) had difficulty scores above 0.5, with an overall average difficulty of 0.33, compared to 15 out of 24 items on the post-assessment (63%), which had an overall average difficulty of 0.57 (Table 1). Similar trends were observed in 2017, with 6 out of 30 items above 0.5 on the pre-test (20%) with an average score of 0.36, compared to 25 out of 30 items on the post-test (83%), which had an overall average difficulty of 0.66 (Table 2). Several questions had difficulty scores that decreased, including items 4, 19, and 24 in 2016 and items 4 and 10 in 2017 (Tables 1 and 2). These items should be revised in future assessments.

**Table 1.** Item Statistics for DFB 2016 - Puerto Rico.

| Concept | Item # | Difficulty<br>Pre | Difficulty<br>Post | Discrimination<br>Pre | Discrimination<br>Post | Bloom's Level |
| --- | --- | --- | --- | --- | --- | --- |
| Early Embryo | 1 | 0.33 | 0.71 | 0.29 | 0.14 | Knowledge |
|  | 2 | 0.54 | 0.83 | 0.14 | 0.14 | Knowledge |
|  | 3 | 0.08 | 0.71 | 0.29 | 0.43 | Comprehension |
|  | 4 | 0.38 | 0.33 | 0.57 | 0.57 | Analysis |
| Gene Expression | 5 | 0.38 | 0.38 | 0.00 | 0.43 | Knowledge |
|  | 6 | 0.42 | 0.50 | 0.00 | 0.29 | Knowledge |
|  | 7 | 0.83 | 0.83 | 0.14 | 0.14 | Knowledge |
|  | 8 | 0.29 | 0.63 | 0.29 | 0.29 | Analysis |
| Cell Signaling | 9 | 0.25 | 0.25 | 0.43 | 0.29 | Knowledge |
|  | 10 | 0.17 | 0.54 | 0.29 | 0.29 | Knowledge |
|  | 11 | 0.58 | 0.83 | 0.00 | 0.14 | Knowledge |
|  | 12 | 0.08 | 0.13 | 0.00 | 0.14 | Application |
| Organogenesis | 13 | 0.38 | 0.42 | 0.00 | 0.29 | Knowledge |
|  | 14 | 0.17 | 0.63 | 0.00 | -0.14 | Knowledge |
|  | 15 | 0.04 | 0.63 | -0.14 | 0.71 | Knowledge |
|  | 16 | 0.17 | 0.67 | 0.14 | 0.86 | Comprehension |
| Disease | 17 | 0.25 | 0.71 | 0.00 | 0.57 | Comprehension |
|  | 18 | 0.25 | 0.42 | -0.29 | 0.71 | Knowledge |
|  | 19 | 0.54 | 0.38 | 0.00 | 0.71 | Knowledge |
|  | 20 | 0.54 | 0.79 | 0.57 | 0.43 | Application |
| Techniques | 21 | 0.29 | 0.58 | 0.57 | 0.43 | Knowledge |
|  | 22 | 0.38 | 0.88 | 0.71 | 0.29 | Knowledge |
|  | 23 | 0.13 | 0.46 | -0.43 | 0.29 | Knowledge |
|  | 24 | 0.42 | 0.33 | 0.29 | 0.43 | Application |
| Mean |  | 0.33 ± 0.19 | 0.57 ± 0.21 | 0.16 ± 0.28 | 0.37 ± 0.23 |  |
|  |  | difficulty < 0.5 |  | discrimination < 0.2 |  |  |
|  |  | difficulty > 0.5 |  | discrimination > 0.2 |  |  |

**Table 2.**
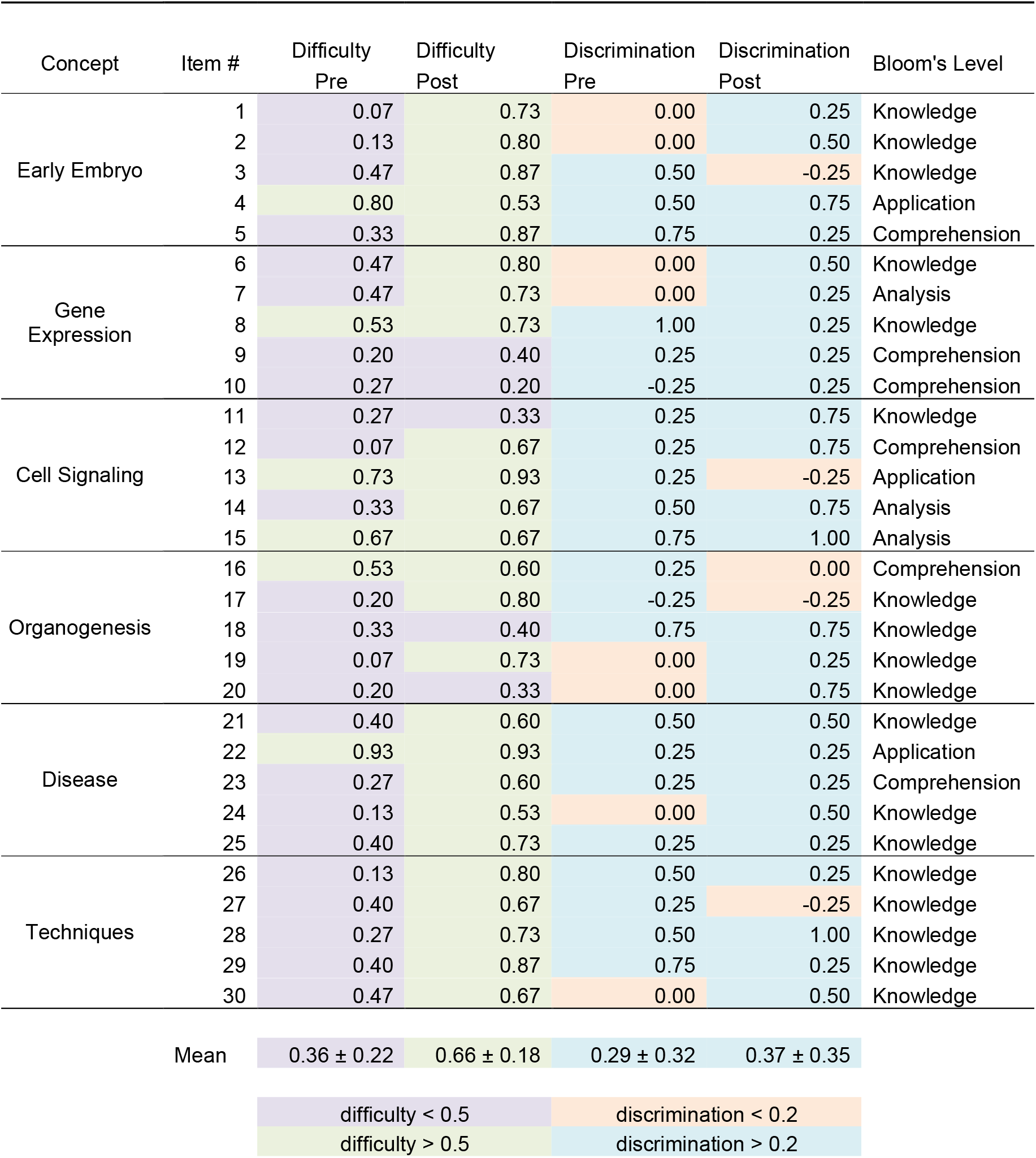
Item Statistics for DFB 2017 - Michigan.

| Concept | Item # | Difficulty<br>Pre | Difficulty<br>Post | Discrimination<br>Pre | Discrimination<br>Post | Bloom's Level |
| --- | --- | --- | --- | --- | --- | --- |
| Early Embryo | 1 | 0.07 | 0.73 | 0.00 | 0.25 | Knowledge |
|  | 2 | 0.13 | 0.80 | 0.00 | 0.50 | Knowledge |
|  | 3 | 0.47 | 0.87 | 0.50 | -0.25 | Knowledge |
|  | 4 | 0.80 | 0.53 | 0.50 | 0.75 | Application |
|  | 5 | 0.33 | 0.87 | 0.75 | 0.25 | Comprehension |
| Gene Expression | 6 | 0.47 | 0.80 | 0.00 | 0.50 | Knowledge |
|  | 7 | 0.47 | 0.73 | 0.00 | 0.25 | Analysis |
|  | 8 | 0.53 | 0.73 | 1.00 | 0.25 | Knowledge |
|  | 9 | 0.20 | 0.40 | 0.25 | 0.25 | Comprehension |
|  | 10 | 0.27 | 0.20 | -0.25 | 0.25 | Comprehension |
| Cell Signaling | 11 | 0.27 | 0.33 | 0.25 | 0.75 | Knowledge |
|  | 12 | 0.07 | 0.67 | 0.25 | 0.75 | Comprehension |
|  | 13 | 0.73 | 0.93 | 0.25 | -0.25 | Application |
|  | 14 | 0.33 | 0.67 | 0.50 | 0.75 | Analysis |
|  | 15 | 0.67 | 0.67 | 0.75 | 1.00 | Analysis |
| Organogenesis | 16 | 0.53 | 0.60 | 0.25 | 0.00 | Comprehension |
|  | 17 | 0.20 | 0.80 | -0.25 | -0.25 | Knowledge |
|  | 18 | 0.33 | 0.40 | 0.75 | 0.75 | Knowledge |
|  | 19 | 0.07 | 0.73 | 0.00 | 0.25 | Knowledge |
|  | 20 | 0.20 | 0.33 | 0.00 | 0.75 | Knowledge |
| Disease | 21 | 0.40 | 0.60 | 0.50 | 0.50 | Knowledge |
|  | 22 | 0.93 | 0.93 | 0.25 | 0.25 | Application |
|  | 23 | 0.27 | 0.60 | 0.25 | 0.25 | Comprehension |
|  | 24 | 0.13 | 0.53 | 0.00 | 0.50 | Knowledge |
|  | 25 | 0.40 | 0.73 | 0.25 | 0.25 | Knowledge |
| Techniques | 26 | 0.13 | 0.80 | 0.50 | 0.25 | Knowledge |
|  | 27 | 0.40 | 0.67 | 0.25 | -0.25 | Knowledge |
|  | 28 | 0.27 | 0.73 | 0.50 | 1.00 | Knowledge |
|  | 29 | 0.40 | 0.87 | 0.75 | 0.25 | Knowledge |
|  | 30 | 0.47 | 0.67 | 0.00 | 0.50 | Knowledge |
| Mean |  | 0.36 ± 0.22 | 0.66 ± 0.18 | 0.29 ± 0.32 | 0.37 ± 0.35 |  |
|  |  | difficulty < 0.5 |  | discrimination < 0.2 |  |  |
|  |  | difficulty > 0.5 |  | discrimination > 0.2 |  |  |

In addition, item discrimination was used to measure how well each assessment question distinguished between high- and low-performing students (Allen & Yen, 2002). Discrimination scores below 0.2 reflected the need for item revision. Interestingly, in 2016, the pre-test had 10 out of 24 questions (42%) adequately discriminating between low and high performing students, while the post-test displayed an increase in discrimination scores, with 18 out of 24 items (75%) above 0.2 (Table 1). Mean discrimination improved from 0.16 on the pre-assessment to 0.37 on the post assessment (Table 1). In 2017, however, both the pre- and post-assessments contained more questions with discrimination scores above 0.2: 22 out of 30 items on the pre-assessment (73%) and 25 out of 30 questions on the post-assessment (83%) (Table 2). In addition, mean discrimination improved for the 2017 pre-assessment (from 0.16 to 0.29), whereas the post-test discrimination mean remained unchanged at 0.37 (Table 2, c.f. Table 1). Six items in 2016 showed decreased discrimination from pre to post (1, 9, 14, 20, 21, and 22), whereas eight items decreased in 2017 (3, 5, 8, 13, 16, 26, 27, and 29). Any item with a discrimination score below 0.2 on the post-test should be considered for revision. Together, these item statistics will guide improvements to the pre-post assessment for future iterations of the course.

Lastly, we used the Blooming Biology Tool to categorize assessment items depending on required cognitive domains (Crowe, Dirks, & Wenderoth, 2008). Because the course was only a week in length, most items measured lower order cognitive skills at the knowledge level (Tables 1, 2). However, we did slightly improve the taxonomy of the test in 2017, with the amount of higher order items increasing from 33% to 40% of the assessment (c.f. Tables 1,2).

### Student experience, but not interest in developmental biology improved after DFB

Because our ultimate goal is to lower barriers to graduate education, we wanted to assess whether students felt they gained experience as a result of the course, and whether the course influenced their intended career pathway in some way. Student perspectives in these areas were assessed via five items in the pre-post analysis (Figure S4, questions 1-5). Over the duration of both the 2016 and 2017 course, students felt they became more familiar with career options in the sciences (Figure S4, Q2. Figure 5, p<0.02). Of note in 2016, the average pre to post test score in this area increased 75% in raw point value (2.88 to 3.36 out of 4). Students also felt that they had gained developmental biology experience in the classroom and the lab in both the Puerto Rico and Michigan courses (Figure S4 Q4 and Q5. Figure 5, p< 0.01). Surprisingly, overall interest in developmental biology did not increase either year (Figure S4 Q1, Figure 5, p≥0.4). These data may reveal a selection bias for students with a strong interest in developmental biology prior to their participation in DFB. Also of interest, the likelihood of students attending graduate school did not change (Figure S4 Q3, Figure 5, p≥0.2). Together, these results indicate that the DFB course improved students’ perceived experience in developmental biology, but not their immediate likelihood of attending graduate school.

**Figure 5.**
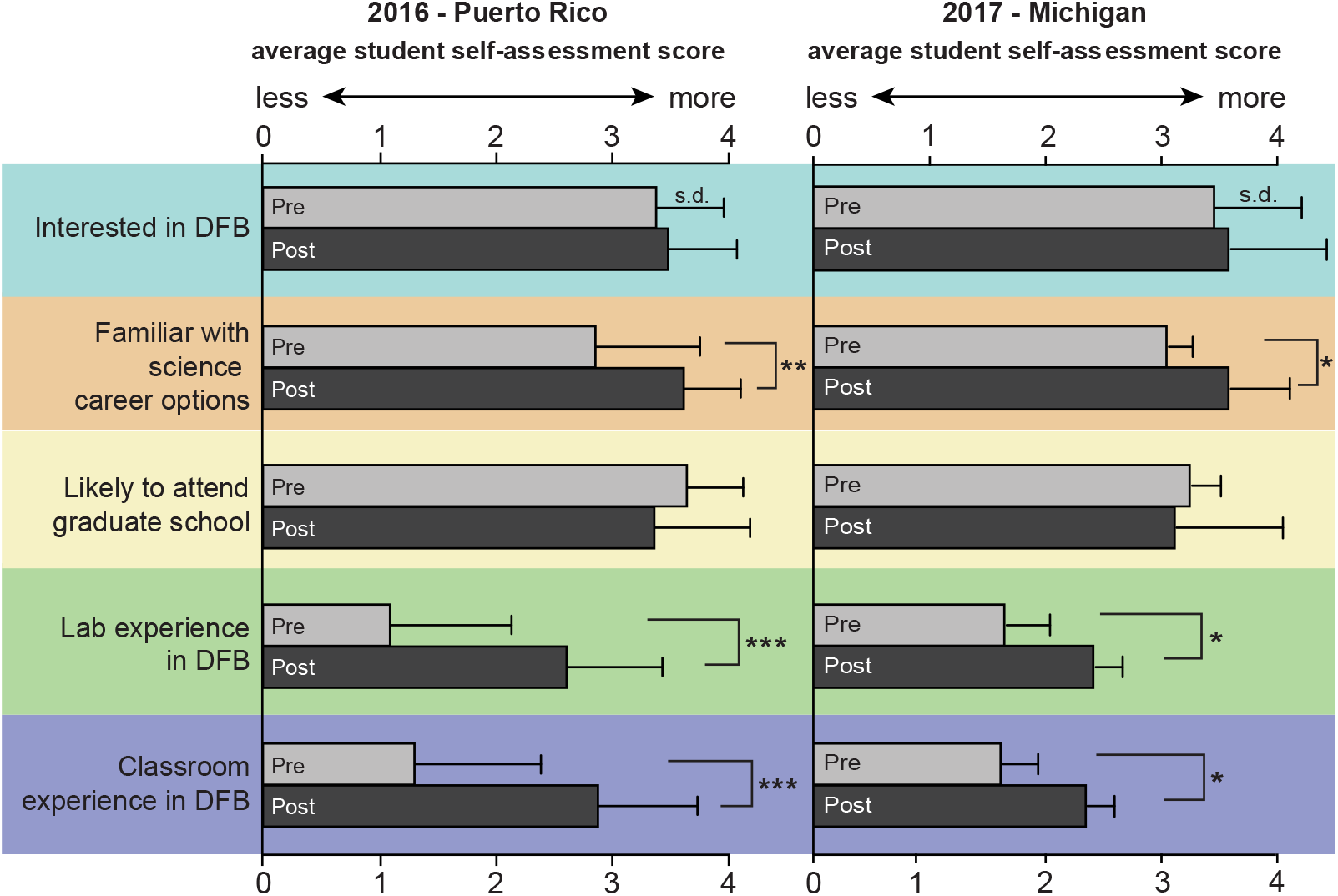
Student attitudes about developmental biology and careers in science. Pre (top, lighter bars) and post (bottom, darker bars) test analysis sorted by item from 2016 (left) and 2017 (right). Student reflection on familiarity with career options in the sciences and experience gained in the classroom and lab significantly improved (career options: p<0.02, and lab and classroom: p< 0.01). Overall interest in developmental biology and likelihood to attend graduate school did not significantly change (p≥0.2) *p<0.02, **p<001, ***p<0.0001.

## Discussion

In this report, we describe the development, implementation, and assessment of Developing Future Biologists, a portable short course in developmental biology. Our mission centers around engaging underrepresented undergraduate students in an active learning environment, while also providing professional development and continued mentorship. Two of our major goals during the course were to teach students the core concepts of developmental biology and to increase awareness of career options in the sciences. Using a pre-post method of assessment, we found that student understanding of core concepts in developmental biology improved, particularly in content areas of organogenesis and early embryonic development. Item analysis indicated that students were able to perform better on post-tests and that discrimination and taxonomy improved with the revised test given for the second iteration of the course. Furthermore, students indicated an increased awareness of career options in the sciences, although their interest in developmental biology and likelihood of attending graduate school did not change. Overall, this course represents a novel educational model that could be widely adopted by other universities and departments to help reduce barriers to graduate education.

### Portable Course Design and the Alumni Connection

Many universities and research institutions offer paid summer research opportunities or internship programs that allow students to gain experience in developmental biology laboratories. For students to pursue these opportunities, however, they must first be motivated to seek out and apply for these programs, and they also must be willing to travel long distances and commit considerable time to these programs (typically around 10 weeks). Arguably, students with little background in developmental biology and with limited access to laboratory resources might be unaware of these summer research programs. Additionally, students may be hesitant to commit an extensive period of time to such endeavors without knowing if they are truly interested in the field. Our unique course model allows us to engage students who might be curious about developmental biology but are unable to commit to a full summer research opportunity in the field. By bringing this course directly to the students, we lower the activation energy required for participation. For students who are interested in pursuing additional research opportunities after the course, our professional development sessions and long-term mentoring model allows us to help them identify subsequent summer research programs and prepare successful applications.

The initial geographical location for the course arose naturally from our instructor alumni connections to UPR Ponce. Not only did our instructors from Ponce recognize the need for this type of initiative based on their own experiences, but they also were able to help tremendously with the course logistics. Given the portable nature of the course, great care was taken to ensure that laboratory activities were feasible with the equipment and resources available at Ponce. Thus, it was incredibly helpful to have alumni involved who were familiar with the institution and facilities. Our alumni instructors also facilitated networking connections between the DFB team and Ponce faculty members, who helped with advertising, coordinating space, and receiving material shipments (all tissues samples used in the labs were shipped to Ponce, with the exception of the locally obtained chicken eggs). Their experience and institutional knowledge was invaluable to the success of the course.

Many groups have demonstrated that shared social identities, including visual identity, have a profound impact on students’ perceptions of themselves in a certain career path (Whittaker & Montgomery, 2012). Our Ponce alumni instructors as well as team members from other UPR campuses enabled course participants to instantly build connections on the basis of shared identity and common experience. DFB participants are therefore able to receive mentoring from positive role models from similar backgrounds to their own, who have already successfully navigated the path to graduate school and a career in science. Overall, we found that developing a partnership with the UPR Ponce community through alumni connections was crucial to the success of the program, and strongly advise that others trying to develop similar programs take this into consideration.

### Course Impact on Participants and Instructors

Although a weeklong course seems like a very short period of time, the outcomes we present here strongly suggest the potential for a lasting impact, both on the UPR students participating in the course and the instructors from the University of Michigan. Within a single week, participants improved their understanding of the core concepts of developmental biology and became more aware of career options in the sciences. While the course was focused on developmental biology, the laboratories and lecture materials exposed UPR and Michigan students to a wide variety of model organisms and research techniques used broadly throughout many fields in biology and biomedical science. Overall, we provided students with a basic introduction to developmental biology, highlighted some of the exciting ongoing research in the field, and helped students more easily envision career paths for themselves in STEM.

Our pre-post assessment indicated that the application process selected for students who were highly interested in developmental biology prior to formal instruction, and that student interest remained high throughout the course of the week. While the course did not increase students’ interest in developmental biology, it did allow interested students to access this topic in a meaningful way, as evidenced by significant increases in participants’ level of developmental biology experience in both the laboratory and the classroom. While these initial assessments show promising results, future courses could largely expand and improve assessment beyond the pre-post test mechanism. Several published indices exist to measure scientific integration (Estrada, 2009) as well as scientific self-efficacy (Chemers, 2006). These and other tools will allow us to improve our understanding of student attitudes and performance in the future.

We observed that DFB did not improve the item score pertaining to likelihood of DFB student to attend graduate school. One complication of this metric was that our students had varying ideas of the definition of graduate school, some which included medical and veterinarian school, and others which did not. Additionally, a short term educational initiative that is only a week in length may not be long enough to change undergraduate students’ career plans. Perhaps a more appropriate question for future surveys might be whether students would be interested in attending a longer course or summer research program in developmental biology. Students were able to have honest conversations with graduate students, postdoctoral fellows, and faculty members to help them make more informed decisions about their career aspirations in the future. For students who do decide to pursue additional career development in the sciences, the extended mentorship aspect of the initiative facilitates access to additional resources and continued support, including letters of recommendation, personal advice, and assistance with applying to research programs. The mentorship model also provides a mechanism to follow up with DFB participants and observe which careers they choose to pursue in the long-term. Continuing assessment will be crucial for past and future DFB courses to analyze impact on recruitment and retention of underrepresented students in scientific fields.

In addition to the impact of the course on enrolled students, DFB also had a profound impact on the team that created, planned, and implemented the course. Few opportunities exist for graduate students to create original learning modules and laboratory activities. This experience has also been invaluable for the graduate students and postdoctoral fellows involved, allowing instructors to develop exceptional teaching, communication, and organizational skills. University of Michigan instructors involved with DFB have a unique opportunity to learn more about students from diverse backgrounds and to better serve in mentoring capacities for these students. Presumably, this will make instructors more comfortable and more likely to engage in cross-cultural mentoring in the future, which will be critical for expanding diversity in the sciences overall. Future iterations of DFB should focus on improving instructors’ mentorship skills through resources like the National Research Mentoring Network (NRMN) and other current initiatives.

### Course Design and Scalability

The DFB initiative provides a framework for the development of similar courses with flexible content and length. At the heart of this program are hands-on, active learning approaches and relationship-building between undergraduate students and their near-peer counterparts (Ramani, Gruppen, & Kachur, 2006). Information gathered from instructor evaluations suggests that these approaches were highly valued by our students. Our assessment data suggest that a week of interactive learning is a sufficient amount of time to teach students core concepts of developmental biology. Our hope is that graduate students, postdocs, and faculty members from other departments and institutions will adapt this model to create similar courses in areas of need. To meet these ends, we formatted the course to be modular in nature so that it could easily be applied in any field of science education.

### Conclusion

To improve diversity outcomes in the sciences, underrepresented students need to be engaged in a meaningful way and shown pathways to success. The weeklong DFB initiative addresses these issues through active learning strategies that have been previously shown to reduce the gap between students of different educational backgrounds (Haak et al., 2011) and by providing mentorship from individuals who have faced similar disadvantages and succeeded, a mentoring strategy demonstrated to increase retention of minority groups in other STEM fields (Dennehy & Dasgupta, 2017; Keller, Logan, Lindwall, & Beals, 2017). Through these methods, DFB was able to build on personal connections, leverage existing diversity, and provide high quality long-term mentoring. We believe these aspects are crucial to the success of DFB and, more importantly, the improvement of the cultural climate in science as a whole.

## Acknowledgements

We are incredibly grateful to a number of individuals, whose contributions were crucial to the success of this course. The Biology department at UPR Ponce was instrumental in planning and implementing this course, providing laboratory space, equipment, and materials as needed. In particular, we thank the administrative personnel of the UPR Ponce: Maritza Torres and Delia Martínez; the professors Heidi Reyes, Dr. Wilfredo Ayala, Dr. Cynthia Rivera, David Forestier, and Dr. Abigail Ruiz; and our invited speakers Dr. Pedro Santiago and Dr. Jose García Arrarás. At the University of Michigan, we thank members of the Allen, Barolo, Spence, Denver, Miller, Innis, and Wellik labs for providing helpful reagents and materials. We also thank Dr. Leah Bricker, Dr. Ken Cadigan, and Dr. Larry Gruppen for helping with course and assessment design, and Dr. Sue Moenter for helping edit the manuscript.

This initiative would also not have been possible without financial support from the Society for Developmental Biology (SDB) Non-SDB Educational Activities Program, The American Society for Cell Biology Committee for Postdocs and Students Outreach, and the following University of Michigan based funding sources: The Rackham Graduate School Dean’s Strategic Fund, the Department of Cell and Developmental Biology, the Endowment for Basic Sciences, the Department of Molecular Cellular and Developmental Biology, the Cellular and Molecular Biology Program, the Department of Molecular and Integrative Physiology, the Center for Organogenesis, the Postbac Research Education Program, the Program in Biomedical Sciences, the Office for Health Equity and Inclusion, the Center for Graduate and Postdoctoral studies, and the generous donors who contributed to our “Giving Blue Day” fundraising campaign.

Importantly, we wish to recognize and thank Martha’s, Andrea’s, Jorge’s, and Leilani’s families, who welcomed us into their homes and helped with various tasks related to the course during the 2015 and 2016 Ponce courses.

Lastly, we would like to thank all of the amazing students who took time out of their summers to enroll in this course and showed up every day full of excitement and curiosity.

